# PymolFold: A PyMOL Plugin for API-driven Structure Prediction and Quality Assessment

**DOI:** 10.1101/2025.10.03.680230

**Authors:** Yifan Deng, Jinyuan Sun

## Abstract

Deep learning has transformed protein structure prediction, yet many experimental scientists face barriers in accessing state-of-the-art (SOTA) models due to technical complexity and hardware requirements. To address this, we present **PymolFold**, an open-source PyMOL plugin that seamlessly integrates cutting edge API-based protein structure predictors such as ESM-3 and Boltz2 into the molecular visualization environment. PymolFold supports both graphical and command-line interfaces for flexible usage and incorporates PXMeter, an open-source Python package for quantitative evaluation of protein structure predictions against reference data. Together, these features establish a unified “predict–visualize–analyze” workflow, lowering technical entry barriers and broadening access to advanced structural modeling. PymolFold is freely available at https://github.com/jinyuansun/PymolFold.

## 1 Introduction

The accurate prediction of three-dimensional (3D) protein structures from amino acid sequences has been a long-standing challenge in molecular biology. Recent breakthroughs in deep learning have transformed this task, with methods such as AlphaFold3, ^1^ RoseTTAFold All-Atom,^2^ Boltz-1/2,^3,4^ Chai-1/2,^5,6^ and Protenix^7^ achieving unprecedented accuracy for both monomeric and complex structures. More recently, language model–based approaches such as ESM-3^8^ have further reduced reliance on multiple sequence alignments (MSAs), accelerating structure prediction workflows.

In parallel, technology companies such as NVIDIA and WeMol^9,10^ are democratizing access to SOTA AI-for-Science models through cloud-based APIs, which eliminates the need for local deployment. Nevertheless, these services remain largely geared toward technically proficient users. Even widely accessible platforms such as the AlphaFold3 server or NVIDIA’s Boltz-2 portal still require repeated transitions between sequence submission, result download, and external visualization in tools like PyMOL.^11^ This fragmentation limits efficient hypothesis testing, while advanced analyses often require additional software. At the same time, emerging interfaces such as ChatMol^12^ highlight the potential of large language models (LLMs) to guide structural studies, but their outputs can be unreliable, with hallucinations or incomplete execution reducing their robustness for research use.

To overcome these barriers, we developed **PymolFold**, a PyMOL plugin that integrates structure prediction, visualization, and evaluation in a single environment. PymolFold leverages multiple prediction models through API access, enabling users to run both monomeric (ESM-3) and multimeric (Boltz-2) predictions without specialized hardware (Figure 1A). Crucially, it incorporates an adapted version of PXMeter^13^ for immediate quantitative benchmarking against reference structures. By consolidating the *predict–visualize–analyze* workflow, PymolFold lowers the technical barrier for computational biology, empowering scientists to generate, inspect, and validate structural hypotheses directly within their preferred visualization environment. The main commands and their functions are summarized in Table 1.

**Table 1:** New Commands Introduced in PymolFold.

| Command | Description |
| --- | --- |
| `esm3 <seq>, <name>` | Predicts a monomer structure from an amino acid sequence. |
| `esmfold <seq>, <name>` | Uses ESMFold to predict the structure of a monomeric protein from an amino acid sequence. |
| `boltz2` | Launches the graphical user interface (GUI) for complex prediction. |
| `color_plddt` | Colors a predicted structure by its pLDDT confidence score. |
| `pxmeter_align <ref>, <model>` | Quantitatively assesses a model against a reference structure. |

**Figure 1.**
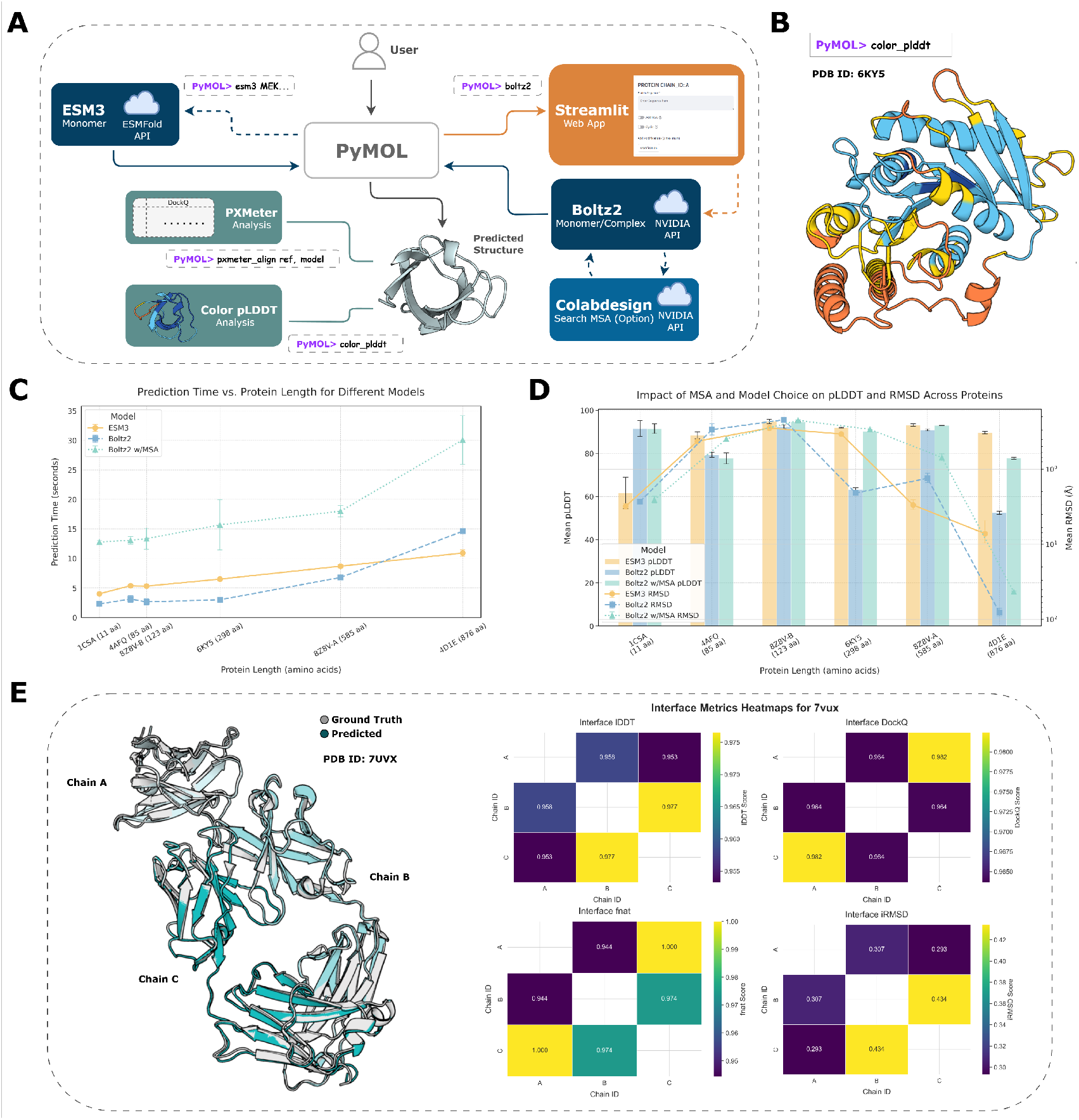
PymolFold workflow, performance case studies, and application to complex structure prediction. **(A)** The PymolFold workflow. A Streamlit GUI in PyMOL calls remote prediction APIs (ESM-3, Boltz2; dashed lines). Results are returned for local analysis with integrated tools (PXMeter, color_plddt; solid lines). **(B)** A representative monomer prediction of a hydrolase mutant (PDB: 6KY5), utilizing color_plddt command, where deep blue indicates high confidence. **(C)** Case studies of prediction time as a function of protein length. **(D)** Performance comparison showing mean pLDDT (confidence) and C*α*-RMSD (accuracy).For (C) and (D), data are the mean ± s.d. of three replicates. **(E)** Predicted trimer (cyan; PDB: 7UVX) superimposed on its ground truth structure (gray). Adjacent PXMeter heatmaps display interface quality metrics (lDDT, DockQ, Fnat, iRMSD).

## 2 Features and Application Workflows

PymolFold is designed to be intuitive, with its core functionalities accessible through simple PyMOL commands. The architecture ensures that PyMOL remains responsive while handling background tasks like API calls and analysis.

### 2.1 Monomer Prediction and Flexible Workflows

To empirically characterize the performance trade-offs between these methods, we conducted case studies across six proteins of varying lengths and origins. The test set was chosen to represent a diverse range of challenges sourced from the RCSB Protein Data Bank: ^14^ the cyclic peptide drug Cyclosporin A (11 amino acids, aa; PDB ID: 1CSA) ; a small engineered scaffold, the Fynomer miniprotein (85 aa; PDB ID: 4AFQ) (Figure 1B); a therapeutic VHH nanobody domain (123 aa; chain B, PDB ID: 8Z8V, 8Z8V_B); a medium-sized globular enzyme (298 aa; PDB ID: 6KY5); the large, multi-domain human serum albumin (585 aa; chain A, PDB ID: 8Z8V, 8Z8V_A); and the very long structural protein alpha-actinin-2 (876 aa; PDB ID: 4D1E).

Our analysis, which includes total time elapsed from API request to response, reveals distinct performance profiles for the three workflows (Figure 1C). For the majority of proteins with lengths under around 600 aa, the MSA-free Boltz2 workflow was consistently the fastest. The automated MSA search adds a notable time overhead, making the Boltz2 with MSA approach the slowest. Interestingly, while ESM-3 was slightly slower than the MSA-free Boltz2 for shorter sequences, its prediction time scaled more favorably with increasing protein length. For the longest protein in our set, alpha-actinin-2 (876 aa), ESM-3 emerged as the faster MSA-free alternative.

In addition to timing, our performance metrics highlight nuanced differences in structural accuracy (Figure 1D). Boltz2, likely benefiting from its specialized cyclic peptide features, achieves high predicted local distance difference test (pLDDT) on 1CSA; however, root mean square deviation (RMSD) shows minimal improvement relative to ESM-3. Across the dataset, incorporation of MSA generally enhances pLDDT for Boltz2, and in proteins such as 6KY5 and 8Z8V_A, RMSD is also significantly reduced, indicating improved structural fidelity. For large multi-domain proteins (4D1E), ESM-3 appears better able to capture overall conformational features, offering a favorable balance of accuracy and computational efficiency.

Taken together, these results provide practical guidance for PymolFold users: for most typical proteins, Boltz2 without MSA offers the fastest and reasonably accurate predictions; for proteins where maximal pLDDT or RMSD improvement is desired, adding MSA to Boltz2 can yield measurable gains; and for very large or multi-domain proteins, ESM-3 represents a competitive, time-efficient alternative. These insights integrate both accuracy and computational considerations, facilitating informed workflow selection.

### 2.2 Complex System Prediction and Validation Workflow

To demonstrate PymolFold’s capabilities for complex systems, we performed a prediction and validation on a therapeutically significant target: the immune checkpoint protein PD-1 bound to a therapeutic Fab antibody fragment (PDB ID: 7VUX). This system is a prime example of a multi-chain protein-protein interaction critical to modern cancer immunotherapy.

The workflow begins in PyMOL with the `boltz2` command, which launches the Streamlit-based GUI. For the 7VUX case, the sequences for the PD-1 extracellular domain (Chain A), and the Fab heavy (Chain B) and light (Chain C) chains were submitted via this GUI, with the automated MSA option enabled.

With the predicted model loaded alongside the fetched crystal structure, the next step is a rigorous, quantitative validation using the `pxmeter_align` command. This function integrates a modified PXMeter engine, which first performs a structural superposition^15^ and then computes a suite of interface quality metrics. The results are exported as both a summary CSV file and a series of intuitive heatmaps (Figure 1E). Key metrics for assessing the predicted protein-protein interfaces include:

- **Interface lDDT:** Assesses the accuracy of the local atomic environment specifically at the protein-protein interface.
- **iRMSD:** The backbone root-mean-square deviation of interface residues after superposition, directly measuring the accuracy of the binding pose.
- **Fnat:** The fraction of native residue-residue contacts successfully recapitulated in the predicted model.
- **DockQ:** A score between 0 and 1 providing a single, continuous measure of docking quality, integrating Fnat and iRMSDs.

The analysis of the 7VUX complex revealed a highly accurate prediction. All pairwise interfaces—PD-1 (Chain A) with Fab heavy chain (Chain B), PD-1 (Chain A) with Fab light chain (Chain C), and heavy-light chain (B-C)—achieved DockQ scores above 0.96. These high scores, together with a low iRMSD (<1.5 Å) and a high Fnat (>0.9), indicate that the predicted model closely matches the native binding mode. The entire process, from multichain sequence input to quantitative interface analysis, was completed in minutes, with this case requiring approximately 2 minutes, highlighting the practical utility of PymolFold for complex system modeling.

## 3 Conclusion

PymolFold provides a powerful and accessible bridge between SOTA structure prediction technologies and the daily workflow of structural biologists. By integrating API-driven prediction for diverse molecular systems and providing an immediate, quantitative assessment module directly within PyMOL, it significantly lowers the barrier to entry and streamlines the validation of structural models. The unified “predict–visualize–analyze” workflow empowers researchers to rapidly iterate on structural hypotheses, accelerating the pace of discovery in structural biology and drug design. Future work will focus on incorporating additional prediction models and expanding the scope of the structural analysis module.

## 4 Methods

### 4.1 Plugin Architecture and Implementation

PymolFold is an open-source Python (v3.9) plugin designed for modern versions of PyMOL (tested on v3.1). Installation is handled by a single run command within PyMOL, which fetches the installer script from the official GitHub repository.

The plugin employs a multi-processed, non-blocking architecture to maintain responsiveness. The `boltz2` command launches a local FastAPI/Uvicorn server in a background thread to handle API requests, while the GUI, implemented as a Streamlit application, runs in a separate `subprocess` . Input validation and molecular data handling are supported by RDKit (v2025.3.6), and analysis and visualization modules leverage Matplotlib (v3.3.0) and Seaborn (v0.13.2).

Structure predictions are performed via external API services, including Meta’s ESM-3 API and NVIDIA’s endpoints for Boltz2 and the ColabDesign MSA server,^16^ with user-provided keys configured as environment variables. This integrated design enables users to perform prediction, visualization, and quantitative analysis seamlessly within the PyMOL environment.

### 4.2 Structural Analysis with PXMeter

The `pxmeter_align` command provides a quantitative assessment of prediction quality. This module integrates a locally modified version of the open-source PXMeter package (v0.1.4), adapted to resolve potential Python dependency conflicts in certain PyMOL sandbox environments and to extend compatibility from Linux-only code to all major operating systems. When executed, the command takes two PyMOL selection strings (a reference and a model) as input, performs a structural alignment, and passes the aligned structures to the PXMeter engine. The engine calculates a comprehensive set of metrics, including global scores (lDDT, RMSD) and interface-specific scores (DockQ, iRMSD, fnat, etc.), which are then saved to a CSV file and rendered as plots.

### 4.3 Availability

The PymolFold plugin, the source code, the installation instructions, and detailed documentation are freely available for academic use on GitHub at https://github.com/jinyuansun/PymolFold.

